# Maturation of lateral habenula and early-life experience-dependent alteration with behavioral disorders in adulthood

**DOI:** 10.1101/2020.04.23.056200

**Authors:** Tomoya Nakamura, Kohei Kurosaki, Munenori Kanemoto, Masakiyo Sasahara, Hiroyuki Ichijo

## Abstract

The lateral habenula (LHb) inhibits midbrain monoaminergic neurons, thereby regulating emotion/cognition. Abnormally high activity in the LHb causes behavioral disorders, but how stressful experiences affect neuronal circuits underlying emotion remains poorly understood. Here, we report the effects of chronic stress on the LHb in postnatal day (P)1-9, P10-20, and P36-45 mice in the pre-, early, and late stages of LHb maturation. At P60, only mice exposed during P10-20 exhibited LHb-specific changes: abnormally high-stress reactivity shown by the expression of the immediate-early gene product (Zif268/Egr1) with insufficient number of parvalbumin (PV) neurons containing GABA. Furthermore, these mice showed anxiety/depression-like behaviors in the light-dark box test/forced swim test. Thus, experiences in early-life are essential for the maturation of neuronal circuits underlying emotion. Early-life stress is thought to have caused anxiety/depression in adulthood by disrupting the maturation of inhibitory PV neurons in the LHb in a period-specific manner.

## Introduction

The lateral habenula (LHb) in the dorsal diencephalon is activated by aversive information from the basal ganglia and the limbic area and subsequently inhibits the activity of downstream monoaminergic systems in the midbrain, such as the ventral tegmental area (VTA) and the dorsal raphe nucleus (DRN), which regulate emotion and cognition (see reviews by Hikosaka et al., 2008; Aizawa et al., 2011; Proulx et al., 2014; Mizumori and Baker., 2017; Yang et al., 2018a). Therefore, it has been suggested that LHb dysfunction causes disorders of behaviors; for example, anxiety and depression.

Anxiety-like behaviors and hyperactivity in the LHb were shown in rats following ethanol withdrawal (Kang et al., 2017). As an animal depression model, learned helplessness has been investigated in detail; the LHb neurons projecting to VTA are activated (Li et al., 2011), the expression of *β*CaMKII is upregulated (Li et al., 2013), the burst firing regulated by the NMDA receptor occurs more frequently (Yang et al., 2018b), and depressive behavior is ameliorated by injection of a GABA agonist into the LHb (Winter et al., 2011). In humans, hyperactivity of LHb neurons has been shown to occur in depression patients (Morris et al., 1999). Thus, it is thought that excessive activity in the LHb is involved in the mechanisms underlying anxiety and depression.

In epidemiological studies, anxiety/depression-related disorders are known to be caused by aversive experiences such as excessive stress during childhood in the form of child neglect and abuse in humans (Li et al., 2016; Rehan et al., 2017). In rodent experiments, anxiety-or depression-like behaviors in adulthood occur after 3 hours of repeated maternal deprivation (RMD) during P1-21 (MacQueen et al., 2003; Aisa et al., 2006). These studies often focused on the hypothalamic-pituitary-adrenal system (HPA), which is a major defense system against stress stimuli; however, involvement of the HPA in behavioral disorders has been differentially reported in studies (see reviews by Millstein and Holmes et al., 2007; Molet et al., 2014). On the other hand, it has been reported that 6 hours of RMD during P7-15 results in insufficient GABA_*B*_R signaling and LHb hyperexcitability, resulting in depression-like behaviors (Tchenio et al., 2017).

Early-life experiences have also been known to disturb the maturation of neurons and to subsequently cause functional and structural neural circuit alterations throughout life, and this property of altering neural circuits is referred to as plasticity. In rodent studies, high neuronal plasticity is reported to depend on high neuronal activity in early-life (Gordon and Stryker., 1996), and in this process, the parvalbumin positive (PV) inhibitory interneurons are involved by suppressing plasticity (Sugiyama et al., 2008; Miyata et al., 2012). Our previous study has shown that the LHb has higher reactivity to stress stimulation in early-life than in adulthood by using the expression of an immediate-early gene (Zif268/Egr1), which is a reliable marker for increased neuronal activity (Ichijo et al., 2015). However, there has been no detailed description of LHb maturation and its relation to plasticity. Therefore, in the LHb, it is worth examining (1) course of maturation in terms of inhibitory neurons, especially PV neurons, to determine the alteration in neuronal activity, (2) experience-dependent plastic changes during maturation, and (3) experience-dependent plastic changes in behaviors.

We examined the maturation of circuits in the mouse LHb, investigated the effects of early-life stress on the LHb, and evaluated how such stress affects lifelong behavior. The stressful experiences in early-life are thought to have caused anxiety and depression in adulthood through disturbing the maturation of the inhibitory GABAergic PV neurons, thus resulting in hyperactivity in the LHb.

## Materials and Methods

### Animals

Wild-type male and female C57BL/6J mice (B6/J) were purchased from Japan SLC Inc. (Hamamatsu, Japan). Heterozygous GAD1 (GAD67)-GFP knock-in mice were provided by RIKEN Bio Resource Center (Tamamaki et al., 2003; RBRC03674, Tsukuba, Japan). To drive the expression of mCherry in GAD2 (GAD65) neurons (GAD2-mCherry mice), GAD2-IRES-Cre female mice (Taniguchi et al., 2011; Stock No. 010802, The Jackson Laboratory, Bar Harbor, USA) were crossed with R26R-H2B-mCherry male mice (Abe et al., 2011; CDB0204K, RIKEN CLST, Wako, Osaka, Japan). Adult mice were maintained in single-sex groups with up to five littermates per cage in a temperature-controlled environment (22-25◦C) with a 12/12-h light/dark cycle (lights were turned on at 5:00, and off at 17:00). Pups were maintained in a cage with a dam. Food and water were supplied ad libitum. Results were similar in cases where males and females were analyzed separately; therefore, we presented pooled data of males and females. All experimental groups were avoided to configure from only one littermate.

### Immunohistochemistry and lectin histochemistry procedure

Under deep anesthesia with pentobarbital (50 mg/kg body weight, i.p.; Kyoritsu Seiyaku, Tokyo, Japan), mice were transcardially perfused with phosphate-buffered saline (PBS) and, subsequently, with 3.7% formaldehyde (Wako, Osaka, Japan) in PBS. Brains were dissected out and kept in the same fixative at 4◦C overnight. After washing in PBS, they were embedded in gelatin (16.7% gelatin and 16.7% glycerol in PBS) and postfixed in the same fixative for 4 days at 4◦C. Coronal sections of 70-*µ*m thickness were cut using a vibratome (VT1000S, Leica Microsystems, Wetzlar, Germany). Alternative sections were collected and stored in PBS containing 0.02% sodium azide at 4°C until staining.

For immuno- or lectin-enzyme-histochemistry, the alternative sections were incubated with primary antibodies in 0.5% Triton X-100 in PBS (PBST) with 5% normal goat serum (16210064, Thermo Fisher Scientific, Waltham, USA; PBSTN) or biotinylated WFA lectin (B-1355, Vector, Burlingame, USA) in PBST for 3 days. The sections were incubated in the appropriate biotinylated secondary antibody for 2 hours: anti-rabbit IgG (1:200; BA-1000, Vector) and anti-mouse IgG (1:200; BA-2000, Vector). After washing, they were incubated for 2 hours in ABC reagent (PK-6100, Vector). The reactive signals were visualized with Metal Enhanced DAB Substrate Kit (34065, Thermo Fisher Scientific). The sections were observed under a microscope (DMRE, Leica), and images were obtained with a digital camera (Ds-Ri1, Nikon, Tokyo, Japan). For double immunofluorescence histochemistry, the alternative sections were incubated for 6 days with primary antibody in 3% bovine serum albumin (9048-46-8, Wako) in PBST. They were incubated with appropriate secondary antibodies coupled with Alexa 488, 594, or 647 (1:200, Invitrogen, Carlsbad, USA; Jackson Immuno Research Labs, West Grove, USA) or streptavidin with Alexa 594 for WFA lectin staining (S11227, Thermo Fisher Scientific), counterstained with 4’,6-diamidino-2-phenylindole (DAPI; 1:10000; D9542, Sigma-Aldrich, St Louis, USA), and mounted on glass slides with Mowiol 4-88 (Merck Millipore, Burlington, USA). Confocal images were obtained with a confocal laser scanning microscope (LSM780, Zeiss, Jena, Germany). The entire length of the LHb, from anterior (at −1.22 from bregma) to posterior (at −2.06 from bregma) (Paxinos and Franklin., 2004), was analyzed. The DG and the BLA including the lateral and basal nuclei were analyzed from anterior (at −1.28 from bregma) to posterior (at −1.70 from bregma). The digital images were analyzed using ImageJ software (National Institutes of Health, Bethesda, MD; http://rwsbweb.nih.gov/ij/).

### GABA immunohistochemistry

STAINperfect Immunostaining Kit was used for GABA immunofluorescence histochemistry (ImmuSmol, Bordeaux, France). Under deep anesthesia, the mice were transcardially perfused with a fixation solution in the kit. Their brains were postfixed in the same fixative for 2.5 hours at 4◦C, embedded in gelatin (16.7%gelatin, 16.7% glycerol in PBS), and postfixed again in 4% paraformaldehyde (26126-25, Nacalai tesque, Kyoto, Japan) in PBS at 4◦ overnight. Coronal sections of 70-*µ*m thickness were cut using a vibratome. Immunofluorescence staining for GABA was performed with the STAINperfect Immunostaining Kit according to the protocol.

### Primary antibodies and lectin

For immuno- or lectin-enzyme-histochemistry, the following primary antibodies or lectin were used: an anti-aggrecan rabbit antibody (1:4500 for DAB staining; 1:500 for fluorescence staining; AB1031, Merck Millipore), an anti-GABA chicken antibody (1:200; IS1036, ImmuSmol), an anti-NeuN mouse monoclonal antibody (1:1500, MAB377, Merck Millipore), an anti-Parvalbumin goat antibody (1:500 for fluorescence staining with NeuN; AB_2571614, FRONTIER INSTITUTE, Sapporo, Japan), an anti-Parvalbumin mouse monoclonal antibody (1:90000 for DAB and fluorescence staining; P3088, Sigma-Aldrich), biotinylated Wisteria Floribunda lectin (1:900 for DAB and fluorescence staining; B-1355, Vector), an anti-Zif268/Egr1 rabbit antibody (1:1500 for DAB staining; 1:300 for fluorescence staining; C-19, Santa Cruz Biotechnology, Dallas, USA), and anti-mCherry animal antibody (1:1000 for fluorescence staining; Novus, Centennial, USA).

### Neuronal activity induced by restraint stress

Mice were physically restrained for 60 min in 50-ml polypropylene centrifuge tubes with 9 air holes of 3-mm diameter (339652, Thermo Fisher Scientific) (single exposure to restraint stress: sRS). After sRS, mice were kept for 60 min in the home cage because the number of Zif268/Egr1 immunopositive cells reached the maximum value in the LHb at 60 min after the stress stimulation (Ichijo et al., 2015). Then, under deep anesthesia, a blood sample was obtained from the left ventricle for plasma corticosterone concentration analysis, and mice were perfusion-fixed, and their brains were processed for immunohistochemistry. Neuronal activity induced by sRS was estimated by the number of Zif268/Egr1 immunopositive cells, which was compared with or without sRS in the LHb and DG. The number of Zif268/Egr1 cells with sRS was not significantly different from that with no sRS in the BLA (S1 Figure).

### Neuronal activity induced by maternal deprivation

Pups were separated from dams and placed in an individual transparent plastic box (L 15 cm D × 21 cm H × 13 cm) with fresh animal bedding at room temperature (22-25◦C) for 2 hours between 10:00 and 12:00 on Postnatal day (P) 9 to P20 (single exposure to maternal deprivation: sMD). After sMD, they were perfusion-fixed immediately, and their brains were processed for immunohistochemistry. Neuronal activity induced by sMD was estimated by the number of Zif268/Egr1 immunopositive cells in the LHb and compared with that under no sMD.

### Application of chronic stresses: repeated maternal deprivation and repeated restraint stress

Mice were treated with repeated maternal deprivation (RMD), in which maternal deprivation was repeatedly performed for 3 hours/day between 8:00 and 12:00 during (1) P1-9 (RMD during the first part of early-life: RMDfp) at 25◦C, or (2) P10-20 (RMD during the second part of early-life: RMDsp) at 22-25°C. The mice were weaned on P30 and thereafter housed with the same sex with normal animal care. Restraint stress was performed for 1 hour/day between 8:00 and 12:00 during P36-45 (Repeated restraint stress: RRS). As a control, mice were not treated with RMD nor RRS. From P60 to P70, the mice were subjected to behavioral experiments or they were perfusion-fixed 1 hour after treatment with or without sRS. Their brains were processed for immunohistochemistry analysis of Zif268/Egr1, PV, and aggrecan, or lectin-histochemistry of WFA.

### Light/dark box test

The light and dark apparatus consisted of two identically sized compartments (L 15 cm × W 15 cm × H 30 cm) connected by a small opening (4 cm × 4 cm). To adjust the brightness of the light compartment to 200 lux, the apparatus was put in a big black box (L 67 cm × D 67 cm × H 97 cm) with five LEDs inside (Y07CLL40W01CHJ, Yazawa, Tokyo, Japan). The apparatus was wiped with 70% ethanol and air-dried before the trials. After placing the mice in the center of the dark compartment, mice in the apparatus were recorded for 5 min with a digital video camera (c920, Logitech, Lausanne, Switzerland). Time spent in the dark compartment was measured.

### Forced swim test

A cylinder (22 cm diameter × 22 cm height) was filled with water up to a height of 15 cm and placed at 25-28◦C in 200 lux. A mouse was placed in the cylinder, and its activities were recorded using an overhead digital video camera for 10 min. Its positions were traced using Motr (Ohayon et al., 2013), which was analyzed with a custom-written MATLAB script. To evaluate its general locomotor activity, the total trace length of its body center was measured. Immobilization was defined as satisfying all of the following conditions. First, acceleration was maintained below 0 for more than 3 sec, which indicated moving at a constant speed, reducing speed, or stopping (Figure 6Ba). Second, speed was maintained at less than 3 cm/s for more than 3 sec; the speed did not drop to 0 by an acting inertial force (Figure 6Bb). Third, change in the distance between the nose and the tail root was less than 0.4 cm/sec for more than 3 sec (Figure 6Bc). Immobilization time was analyzed from 5 to 10 min. After the test, the mouse was dried with a paper towel and returned to the cage. The cylinder was washed with water and dried between trials.

### Plasma corticosterone concentration

After puncturing the left ventricle immediately before perfusion under deep anesthesia with pentobarbital (50 mg/kg body weight, i.p.), blood samples were collected with a 26-gauge needle and syringe washed with heparin (5,000 units/5 mL; Mochida Pharmaceutical, Co., Tokyo, Japan) in an ice-cooled tube. To minimize the influence of circadian fluctuations, the blood samples were collected between 11:30 and 12:30. The samples were centrifuged (3000 g, 10 min, 4◦C), and plasma samples were stored at −80◦C until the assay. The concentration of plasma corticosterone was measured using an AssayMaxTM Corticosterone ELISA kit (Assaypro LLC, Missouri, USA). The absorbance at 450 nm wavelength was read on a microplate reader (Multiskan FC, Thermo Fisher Scientific). Mean values were calculated for triplicate sets of standards and samples. The concentrations of corticosterone were determined from the standard curve.

### Statistical analysis

Data were expressed as means ± SEM. They were analyzed using the Kruskal-Wallis rank-sum test followed by the Steel-Dwass multiple comparison test, simple regression test, Student’s t-test, and one-way analysis of variance (ANOVA) followed by the Tukey HSD test or two-way ANOVA. The statistical significance level was set at *p* < 0.05. All statistical data processing was performed using JMP ® Pro 14.2) (SAS Institute, Cary, USA) or Excel (Microsoft, Redmond, USA).

## Results

### Characterization of PV neurons in the LHb

Although the presence of PV neurons is known in the LHb, their detailed character has not been shown; we scrutinized PV neurons with reliable markers. In the adult LHb of the control mice at postnatal day 60 (P60), 99.9 ± 0.1% of PV cells were immunostained using anti-NeuN antibody; thus, PV cells were determined to be neurons (Figure 1A, B, G). Furthermore, 56.4 ± 6.4% of PV neurons showed high-intensity immunostaining signals of anti-GABA antibody in their cell processes and weak intensity in their cell bodies (Figure 1C, G). However, in glutamate decarboxylase 2 (GAD2)-mCherry mice, only 4.8 ± 0.9% of PV neurons were GAD2-positive in the LHb (Figure 1D, G). In addition, GAD1 cells were not observed in the LHb of GAD1-GFP knock-in mice; all PV neurons were GAD1 negative (Figure 1E, G). Together, these results indicated that PV neurons in the LHb contained GABA without expression of either GAD2 or GAD1. The immunostaining signals of anti-GABA antibody were validated by colocalization with glutamate decarboxylase 2 (GAD2)-mCherry signals in the cerebral cortex and the LHb (S2 Figure).

**Figure 1.**
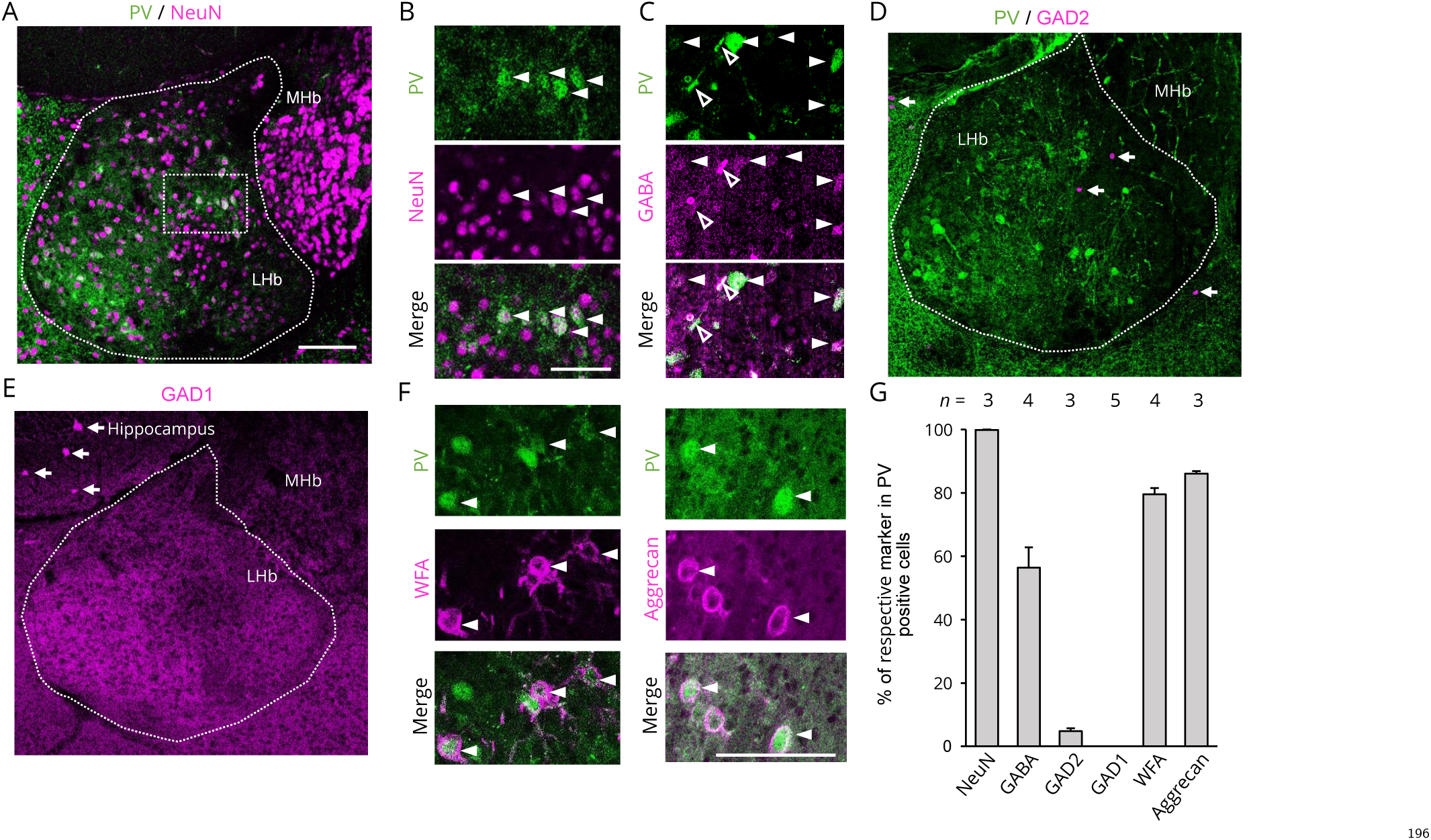
PV neurons in the LHb contain GABA; double immunofluorescence staining of PV, NeuN, GABA, and PNNs. (**A**) Double immunofluorescence staining of PV (green) and NeuN (magenta) is shown in low magnification. (**B**) High-magnification image of the dotted rectangle area in (A) showing PV-positive cells are doubly stained with NeuN (closed arrowheads). (**C**) PV neurons are positive for GABA immunoreactivity on their somas (closed arrowheads) and processes (open arrowheads). (**D**) A few GAD2 reporter-positive cells are found in the LHb (magenta, closed arrows in GAD2-mCherry mice); however, PV neurons (green) are negative for the GAD2 reporter (magenta). (**E**) GAD1 reporter-positive cells are not found in the LHb but seen in the hippocampus (magenta, closed arrows in GAD1-GFP knock-in mice). (**F**) PV neurons (green) are surrounded by PNNs, as determined using WFA lectin staining and aggrecan (magenta). (**G**) Percentage of PV neurons that were positive for the respective markers. Figures A, B, C, and F are photomicrographs of a single optical section. Figures D and E show omnifocal images with 10 optical sections. Closed arrowheads indicate double fluorescence positive cells. Open arrowheads indicate PV and GABA double-positive cell processes. LHb, lateral habenular nucleus; MHb, medial habenular nucleus; PV, parvalbumin; GAD, glutamic acid decarboxylase; WFA, wisteria floribunda agglutinin. The position of the images was near 1.92 mm posterior from bregma. Scale bars = 100 *µ*m.

Most PV neurons in the LHb were surrounded by perineuronal nets (PNNs), which were demonstrated using Wisteria floribunda agglutinin (WFA) lectin and anti-aggrecan antibody. Further, 79.6 ± 1.9% of the PV neurons were positive for WFA lectin staining and 86.1 ± 0.7% for anti-aggrecan immunostaining (Figure 1F, G).

### Maturation of the LHb from early-life to adulthood: PV neurons and PNNs

LHb maturation was examined from early-life to adulthood; the number of PV neurons was counted. In the LHb of the control mice, the number of PV neurons was significantly higher at P60 than at P20 (Figure 2Aa-c). In a simple regression test, the number of PV neurons positively correlated with the number of postnatal days, which indicated that the number of PV neurons in the LHb increased with age (Figure 2Ad).

**Figure 2.**
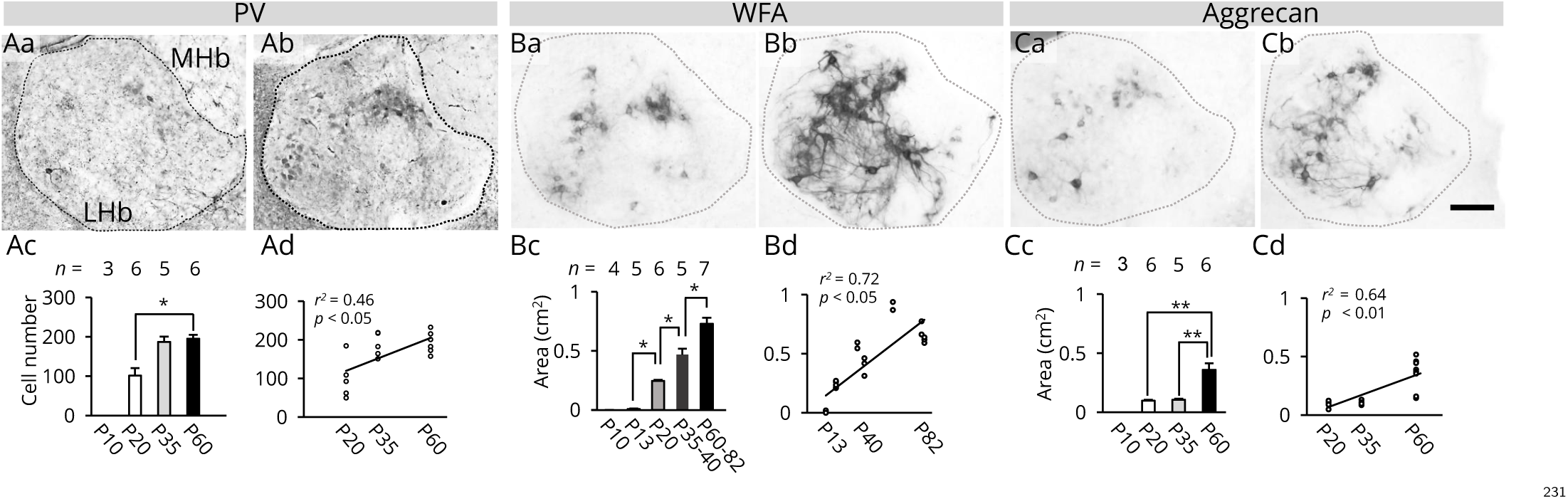
Maturation of PV neurons and PNNs in the LHb. (**A**) PV neurons are shown at P20 (Aa) and P60 (Ab). Their numbers are compared with each other at each postnatal day (*χ*^2^ = 7.99, *p* < 0.05, Kruskal-Wallis rank-sum test) (Ac). The correlation between postnatal day and number of cells is shown using a simple regression test (Ad). (**B**) WFA lectin staining image is shown at P20 (Ba) and P60 (Bb). Their areas are compared at each postnatal day (*χ*^2^ = 20.32, *p* < 0.01, Kruskal-Wallis rank-sum test) (Bc). Correlation between postnatal day and WFA lectin staining area is shown using a simple regression test (Bd). (**C**) Anti-aggrecan immunoreactivity is shown at P20 (Ca) and P60 (Cb). Their areas are compared at each postnatal day (*χ*^2^ = 13.78, *p* < 0.01, Kruskal-Wallis rank-sum test) (Cc). Correlation between postnatal day and anti-aggrecan immunoreactivity area using a simple regression test (Cd). Asterisks and double asterisks indicate significant differences (* *p* < 0.05, ** *p* < 0.01, Steel-Dwass multiple comparison test). Error bars represent SEMs. The square correlation coefficient (r) and p-values of the slope are shown. LHb, lateral habenular nucleus; MHb, medial habenular nucleus; PV, parvalbumin; WFA, wisteria floribunda agglutinin. Scale bar = 100 *µ*m.

The area covered by PNNs was significantly larger in adulthood than during early-life and adolescence; larger at P60-82 than at P35-40, at P35-40 than at P20, and at P20 than at P13, by using WFA (Figure 2Ba-c). Similarly, the area covered by PNNs was larger at P60 than at P20 and P35 (Figure 2Ca-c) as determined using anti-aggrecan. In a simple regression test, the PNN area positively correlated with the number of postnatal days, which indicated that the total area of PNNs increased with age (Figure 2Bd, Cd).

### Maturation of the LHb from early-life to adulthood: its neuronal reactivity during stress

Activation of the LHb was examined after stress stimulation from early-life to adulthood. To determine neuronal activity with no stimulation, the number of Zif268/Egr1 immunopositive cells (Zif268/Egr1 cells) was counted. The number was significantly higher at P20-21, P35, and P61-70 than at P11 (Figure 3Aa-c). To determine neuronal activity in response to stress stimulation during early-life, after a single exposure to maternal deprivation (sMD) as stress stimulation, the number of Zif268/Egr1 cells was significantly higher at P16 and P20 than at P9 (Figure 3Ba-c). In a simple regression test, the number of Zif268/Egr1 cells after sMD positively correlated with the number of postnatal days (Figure 3Bd). To determine neuronal activity in response to stress stimulation after P20, a single exposure to restraint stress stimulation (sRS) was used. In advance, the LHb responses were compared between with and without stimulations at P20; the number of Zif268/Egr1 cells was significantly different between the conditions (control, 83.8 ± 16.4 cells; sMD, 381.8 ± 91.6 cells; sRS, 434.3 ± 94.9 cells; *χ*^2^ = 10.7, *p* < 0.01, Kruskal-Wallis rank-sum test), the number after sMD (*p* < 0.05, Steel-Dwass multiple comparison test) and the number after sRS (*p* < 0.05, Steel-Dwass multiple comparison test) were higher than the number of control. Moreover, the number after sMD is not significantly different from the number after sRS (*p* = 0.99, Steel-Dwass multiple comparison test), indicating the intensities of these stimulations were comparable. In contrast, the number of Zif268/Egr1 cells after sRS was significantly lower at P61-70 than at P20-21 (Figure 3Ca-c). In a simple regression test, the number of Zif268/Egr1 cells tended to negatively correlate with the number of postnatal days during P20-70 (Figure 3Cd). In the adult LHb after sRS, 97.4 ± 8.4% of the Zif268/Egr1 cells were negative for PV immunoreactivity; thus, most cells activated by stress were not PV neurons (Figure 3D). These results showed that neuronal reactivity increased during P10-20 as the mice aged and then became low in adulthood (from P61 to P70).

**Figure 3.**
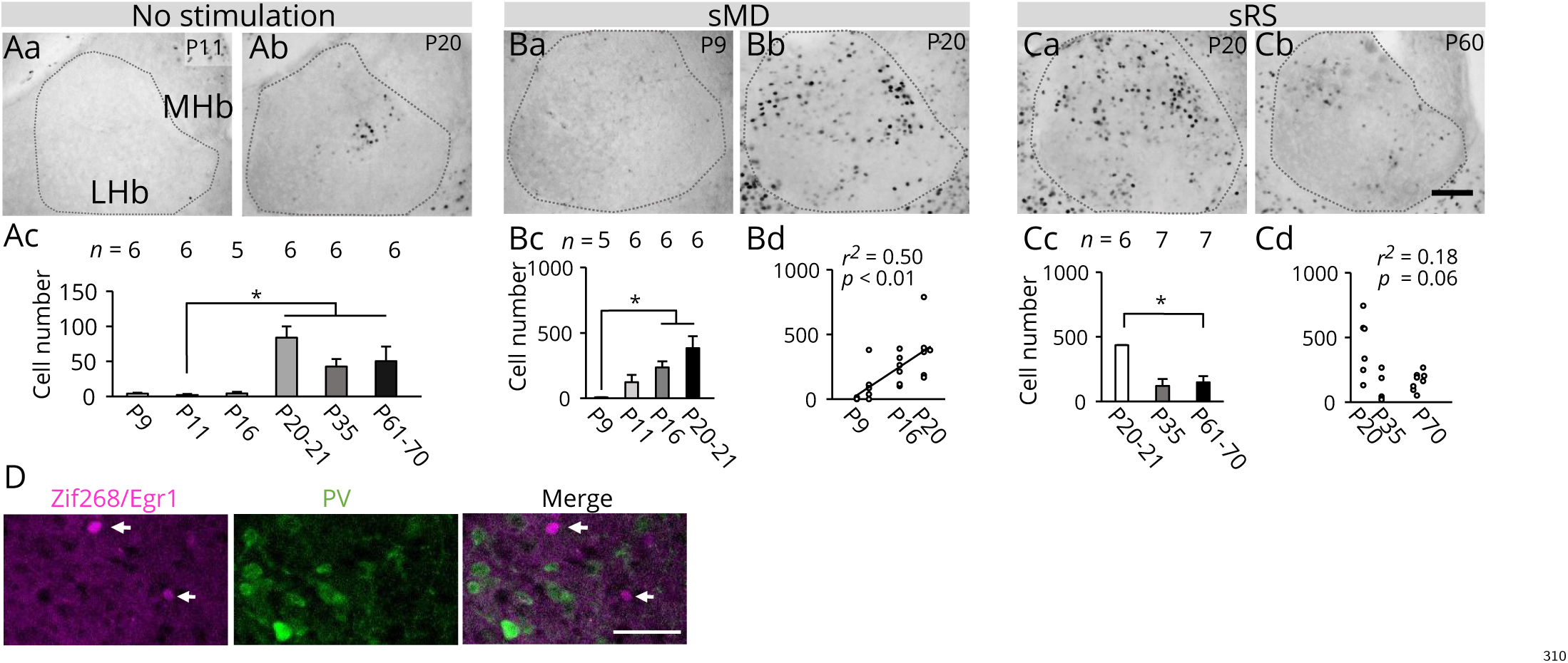
Postnatal changes in neuronal reactivity in response to stress stimulation during the maturation of the LHb. (**A**) Zif268/Egr1-positive cells are shown at P11 (Aa) and P20 (Ab) with no stress stimulation. Their numbers are compared at each postnatal day (*χ*^2^ = 26.27, *p* < 0.01, Kruskal-Wallis rank-sum test) (Ac). (**B**) The positive cells at P9 (Ba) and P20 (Bb) after sMD. Their numbers are compared at each postnatal day (*χ*^2^ = 14.29; *p* < 0.01, Kruskal-Wallis rank-sum test) (Bc). Correlation between postnatal day and number of cells using a simple regression test (Bd). (**C**) Positive cells at P20 (Ca) and P60 (Cb) after sRS. Their numbers are compared at each postnatal day (*χ*^2^ = 7.71, *p* < 0.05, Kruskal-Wallis rank-sum test) (Cc). No correlation between postnatal day and number of positive cells was observed using a simple regression test (Cd). (**D**) LHb was double stained with Zif268/Egr1 (magenta; arrows) and PV (green; open arrows) after the sRS; no cell was double stained. Asterisks indicate significant differences (* *p* < 0.05, Steel-Dwass multiple comparison test). Error bars represent SEMs. The squared correlation coefficient (r) and p-values of the slope are shown. LHb, lateral habenular nucleus; MHb, medial habenular nucleus; sMD, single exposure to maternal deprivation; sRS, single exposure to restraint stress; PV, parvalbumin. Scale bar = 100 *µ*m.

### Alteration in LHb maturation by early-life stress

Maturation of the LHb was examined under the influence of chronic stress during P1-9 (RMD during the first part of early-life: RMDfp), P10-20 (RMD during the second part of early-life: RMDsp), and P36-45 (repeated restraint stress: RRS) (Figure 4A). At P60, significantly fewer PV neurons in the LHb were observed in the RMDsp group than in the control (with no chronic stress), RMDfp, and RRS groups (Figure 4Ba-e). In contrast, in the dentate gyrus (DG) and basolateral amygdala (BLA), the number of PV neurons was not significantly different between the control and RMDsp groups (Figure 4Ca-c, Da-c). In the LHb, the areas positive for WFA and aggrecan staining (PNNs) did not significantly differ between the RMDsp and control groups (S3 Figure).

**Figure 4.**
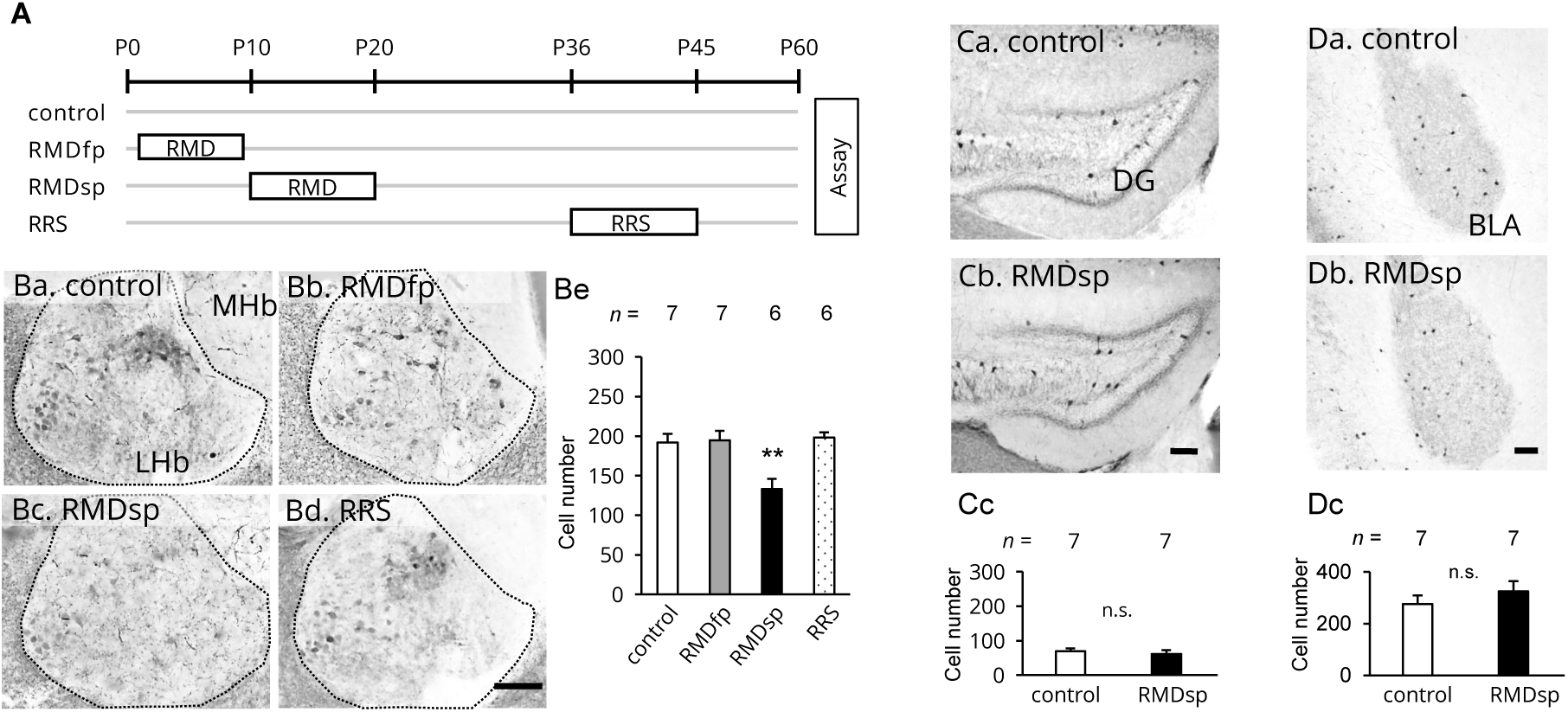
Impaired maturation in the LHb under the influence of early-life stress. (**A**) Schematic of experimental design. RMDfp, repeated maternal deprivation during the first part of early-life (P1-9); RMDsp, repeated maternal deprivation during the second part of early-life (P10-20); RRS, repeated restraint stress (P36-45). (**B**) In the LHb, PV neurons are shown in the control (Ba), RMD during P1-9 (RMDfp) (Bb), RMD during P10-20 (RMDsp) (Bc), and RRS groups (Bd). The number of neurons is compared between groups (F_3,22_ = 9.13, *p* < 0.01, one-way ANOVA) (Be). (**C**) In the DG, PV neurons are shown in the control (Ca) and RMDsp groups (Cb). The number of neurons is compared between the two groups (*p* = 0.57, Student’s t-test) (Cc). (**D**) PV neurons in the BLA (Da, Db). Neuron numbers are compared between groups (*p* = 0.34, Student’s t-test) (Dc). A double asterisk indicates significant difference (** *p* < 0.01, Tukey’s HSD test). Error bars represent SEMs. LHb, lateral habenular nucleus; MHb, medial habenular nucleus; DG, dentate gyrus; BLA, basolateral amygdala; PV, parvalbumin. Scale bar = 100 *µ*m.

Neuronal reactivity in the LHb was evaluated by examining the number of Zif268/Egr1 cells after sRS as the stress stimulation or with no stimulation. In the LHb, the number of Zif268/Egr1 cells after the stimulation was significantly higher in the RMDsp group than in the control, RMDfp, and RRS groups, respectively (Figure 5Aa-e), although the number with no stimulation was not significantly different between the groups (Figure 5Af). However, in the DG, the number after the stimulation was not significantly different between the control and RMDsp groups (Figure 5Ba-c). The plasma corticosterone concentration after stimulation was significantly higher than that with no stimulation, indicating the occurrence of general stress responses (Figure 5C). Furthermore, there were no significant differences in the plasma corticosterone concentration between the control, RMDfp, and RMDsp groups with or without the stimulation, showing that the resting condition and general stress response were similar across groups (Figure 5C). The results showed that the RMDsp had specific effects on the LHb, resulting in fewer PV neurons and higher neuronal reactivity to sRS. Thus, LHb maturation altered under the influence of early-life stress during P10-20.

**Figure 5.**
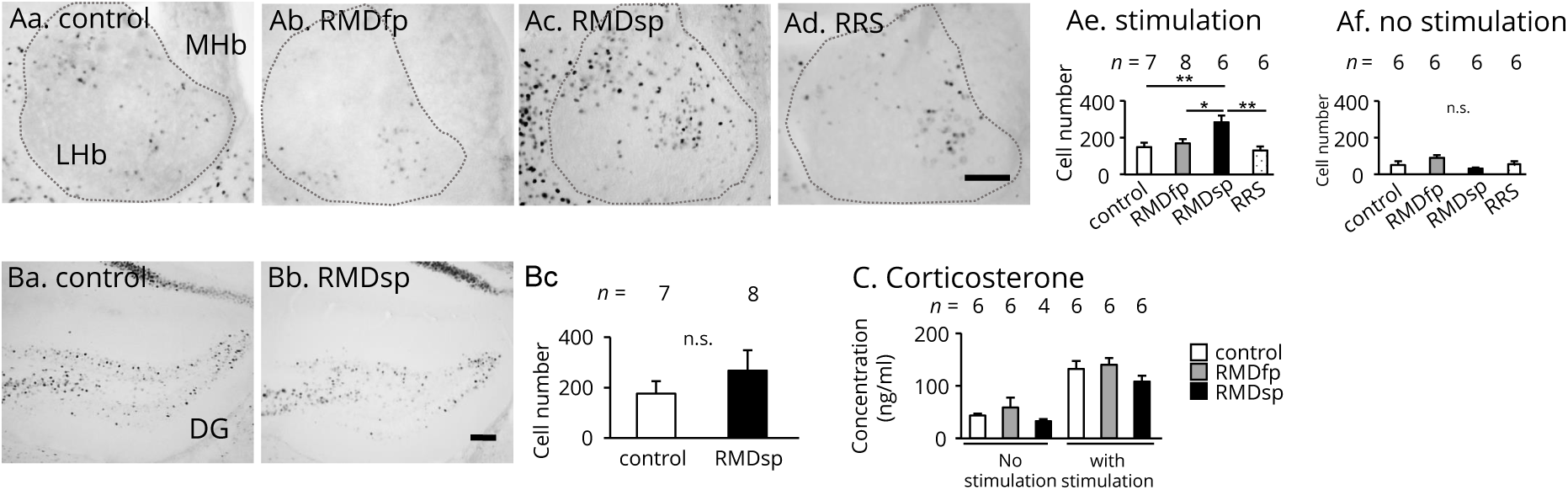
Neuronal hyperreactivity in the adult LHb, with its maturation impaired. (**A**) In the LHb, Zif268/Egr1-positive cells in response to stimulation are shown in the Control (Aa), RMDfp (Ab), RMDsp (Ac), and RRS groups (Ad). Their numbers are compared between the groups with stimulation (F_3,23_ = 6.59; *p* < 0.01, one-way ANOVA) (Ae) and with no stimulation (F_3,20_ = 2.72; *p* = 0.72, one-way ANOVA) (Af). (**B**) In the DG, the positive cells after stimulation are shown in the control (Ba) and RMDsp groups (Bb). Their numbers are compared (*p* = 0.37, Student’s t-test) (Bc). (**C**) Plasma corticosterone concentrations with or without the stimulation are compared between the control, RMDfp, and RMDsp groups (main effect of stress, F_1,28_ = 59.11, *p* < 0.01; main effect of treatment, F_2,28_ = 2.40, *p* = 0.11; interaction, F_2,28_ = 0.12, *p* = 0.88, two-way ANOVA). Asterisk and double asterisks indicate significant differences (* *p* < 0.05, ** *p* < 0.01, Tukey’s HSD test). Error bars represent SEMs. LHb, lateral habenular nucleus; MHb, medial habenular nucleus; DG, dentate gyrus; RMDfp, repeated maternal deprivation during the first part of early-life (P1-9); RMDsp, repeated maternal deprivation during the second part of early-life (P10-20); RRS, repeated restraint stress (P36-45). Scale bar = 100 *µ*m.

### Behavioral disorders under the influence of early-life stress

Influences of early-life stress on behaviors were examined. In the light/dark box test, adult mice in the RMDsp group spent a longer time in the dark compartment than did the mice in the control group (Figure 6A), showing anxiety-like behaviors. In addition, in the forced swim test, the immobilization time was significantly higher in the RMDsp group than in the control group (Figure 6Ca), indicating that the RMDsp mice showed depression-like behaviors. There was no significant difference in the total locomotion distances between the control and RMDsp groups (Figure 6Cb), showing that general locomotor activity was not impaired in the RMDsp group. The results indicated that early-life stress during P10-20 elicited behavioral disorders, anxiety and depression, in adults.

**Figure 6.**
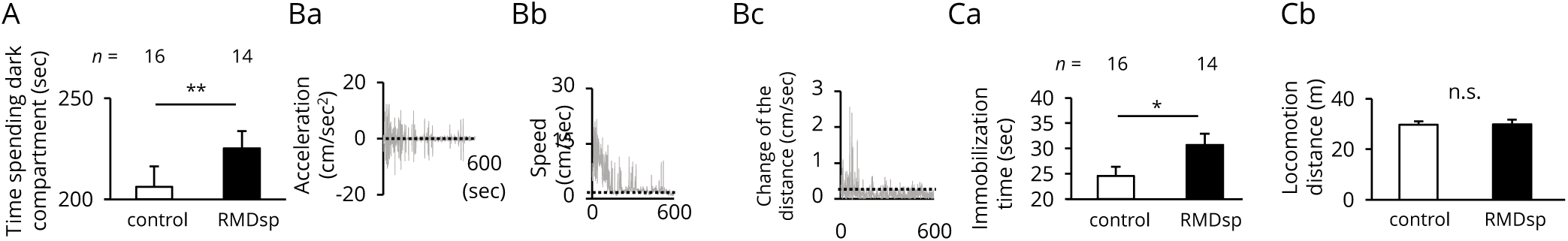
Anxiety/depression-like behaviors in adults exposed to early-life stress. Anxiety/depression-like behaviors in adults exposed to early-life stress. (**A**) In the light/dark box test, time spent in the dark compartment is compared between the control and RMDsp groups. (**B**) Time courses of behavioral activities, acceleration (Ba), speed (Bb), and the distance between the nose and the root of the tail (Bc) are shown for the forced swim test for a control mouse. Immobilization behavior was defined in Material and Methods; dashed lines indicate thresholds of immobilization. (**C**) In the forced swim test, immobilization times (Ca) and locomotion distances (Cb) are compared between the control and RMDsp groups. Asterisk and double asterisk indicate significant differences (* *p* < 0.05, ** *p* < 0.01, Student’s t-test). Error bars represent SEMs. RMDsp, repeated maternal deprivation during the second part of early-life (P10-20).

## Discussion

This study revealed the influence of aversive experiences in early-life on LHb maturation and on behavior during adulthood, indicating experience-dependent mechanisms modulating LHb neuronal circuits. As failure in LHb maturation is likely involved in the etiology of anxiety and depression, this mechanism is thought to be a key to the prevention and treatment of these disorders.

Geisler et al. (2003) reported the presence of PV neurons in the LHb. Although PV neurons are generally known to be inhibitory interneurons, that has not been shown in the LHb. Signals of GABA immunoreactivity were observed in the processes and somas of PV neurons in the LHb (Figure 1C). In addition, they were surrounded by PNNs, similar to the observations in other brain regions (Figure 1F) (Miyata et al., 2012; Yamada et al., 2015; Enwright et al., 2016). Expression of GAD1 mRNA has been shown in the LHb of PV-T2A-Cre mice in the Allen Brain Atlas (https://portal.brain-map.org/); however, there has been no published report on the expression of GADs in PV neurons in the LHb, and genetic reporters for GAD1 and GAD2 were not detected in this study either (Figure 1D, E). Thus, it is likely that PV neurons are GABAergic, expressing a small amount of GAD1 and GAD2 below the detection level of the method used here. In addition, it is also possible that they take up GABA from the extracellular fluid through GABA transporters as previously reported (Pietrini et al., 1994; Minelli et al., 1995; Ribak et al., 1996).

A previous study showed that neuronal activity in the LHb is high in early-life and moderate in adulthood (Ichijo et al., 2015). This study confirmed the former result and unfolds further investigation about LHb maturation (Figure 3). The LHb gradually matures; the four stages are recognized by the extent of neuronal activity, the number of PV neurons, and the area taken up by PNNs. The first stage of LHb maturation is around P10, during which no or low neuronal activity was observed and neither PV neurons nor PNNs were detected (Figures 2 and 3). The LHb is thought to be immature at this stage. The second stage is around P20. During this stage, large spontaneous activity, maximum reactivity against stimuli, and appearances of PV neurons and PNNs were observed (Figures 2 and 3). The LHb is thought to mature in this stage. From the first to the second stage, neuronal reactivity gradually increased in response to stimulation (Figure 3Bc, Bd). The third stage is around P35, during which low spontaneous activity and weaker reactivity against stimulation were observed. The number of PV neurons and the area covered by PNNs increased, indicating that the LHb has functionally matured through the inhibitory PV neurons (Figures 2 and 3). The fourth stage is after P35; during this stage, the area covered by PNNs exhibits additional expansion, indicating further maturation of the LHb (Figure 2B, C). These results show that after P20, the number of PV neurons is inversely related to the neuronal activity of the LHb, indicating that the functional development of PV neurons is involved in LHb maturation. By obtaining information about stages of the LHb maturation in detail, we established experimental set-up for LHb plasticity under emotional experiences in early-life.

It is known that early-life experiences disturb the maturation of neurons and to subsequently cause plasticity: functional and structural neural circuit alterations throughout life. For example, in the visual system, PV interneurons restrict local circuits plasticity under experiences during maturation, which consequently influences higher function (Sugiyama et al., 2008; Miyata et al., 2012, Ribic et al., 2019). Here, we showed PV neurons as neuronal components of plasticity in the LHb under experiences in early-life. In mice subjected to early-life stress at a specific period (during P10-20), there existed fewer PV neurons in the adult LHb with abnormally high neuronal reactivity, following anxiety/depression behaviors (Figures 4, 5, and 6). Early-life stress-induced further late effects on higher brain function in adults, revealing experience-dependent plastic changes in the LHb.

However, since it is not yet known whether the PV neurons in the LHb are interneurons or long projection neurons, there are two hypotheses formulated as mechanisms by which insufficient number of PV neurons in the LHb contributes to the behavioral disorders. In the first hypothesis, as inhibitory interneurons, the PV neurons negatively regulate excitatory long projection neurons in the LHb. Insufficient number of PV neurons leads to hyper-reactivity of the LHb neurons, which exerts effects on inhibitory interneurons in the monoaminergic systems and generate feedforward hyper-inhibition in downstream monoaminergic systems, inducing depression and anxiety by shortage of monoamines (Omelchenko et al., 2009; Brown et al., 2017; Zhou et al., 2017). In the second hypothesis, the PV neurons are assumed to be long projection neurons because PV neurons are identified as an excitatory or inhibitory long projection neurons in outside of the cortex (Kisner et al., 2018; Knowland et al., 2019). As excitatory long projection neurons, the PV neurons positively regulate the monoaminergic system. Insufficient number of PV neurons fails in activating the downstream monoaminergic systems. Or, as inhibitory long projection neurons, the PV neurons regulate inhibitory interneurons in the monoaminergic system. Insufficient number of inhibitory PV projection neurons fails in disinhibiting monoaminergic system, inducing depression and anxiety by shortage of monoamines.

Present study showed that most of the PV neurons were GABAergic (Figure 1C), that fewer neurons responded to stress after PV maturation (Figures 2 and 3), and that insufficient number of PV neurons and neuronal hyper-reactivity were observed in the same early-life stress group which expressed anxiety- and depression-like behaviors (Figures 4, 5, and 6). These results strongly support that the PV neurons are inhibitory interneurons as proposed in the first hypothesis. On the other hand, small number of PV neurons were retrogradely labeled by injection of cholera toxin subunit B in DRN and VTA (data not shown), possibly suggesting that the PV neurons long-project to monoaminergic system. It remains to be elucidated how alteration of LHb neuronal circuits induces anxiety/depression.

The late effects of early-life stress during P10-20 are specific to the LHb spatiotemporally because the early-life stress did not act in the DG and BLA (Figures 4C, D, and 5B) and had no effect on the systemic reaction of HPA late in adults (Figure 5C). Moreover, effective stimulation for inducing LHb plasticity is thought to be restricted in a narrow time window, showing that physically mild stress, specifically in P10-20 (RMD) mice, induces long-lasting influences on behaviors. The stress stimulations during P1-9 (RMD) and P36-45 (RRS) were not effective even though the intensities of these stimuli were thought to be comparable because both sMD and sRS at P20 induced activations with a similar number of Zif268/Egr1-positive cells in the LHb. In contrast, the degree of plasticity in adults is thought to be low. It is reported that intense stress stimuli applied to adult mice have functional effects on the LHb; these stimuli include continuing electrical shocks until learned helplessness (Winter et al., 2011; Li et al., 2011, 2013; Yang et al., 2018b), various stress stimuli (electrical shock, 40 hours fasting, 4◦C swimming, 45◦C heat-stress, 24 hours water-deprivation, 1 min clip tail) (Yang et al., 2008), and 90-180 min restraint stress in a narrow 50-ml tube, which limits forward and backward movement by blocks for 14 days (Yang et al., 2018b; Seo et al., 2018).

Plasticity in early-life is well known as a critical period, which is demonstrated in sensory neuronal circuits. During the critical period, neuronal activity is high, and structure-function relationships are modified by experiences (Gordon and Stryker., 1996). Maturation of circuits reduces plasticity and closes the critical period through the development of inhibitory PV neurons (Sugiyama et al., 2008; Miyata et al., 2012). Here we classified LHb maturation by neuronal components of plasticity. The LHb is highly plastic during the early-life period, in which experiences modify the circuits of PV neurons and cause alteration in late behaviors in adult. LHb plasticity is restricted after the maturation of PV neurons and PNNs. These observations support the notion of a critical period of aversive information processing in the LHb, proposing the hypothesis that the LHb under early-life stress is involved in the etiology of anxiety and depression. The results also support the use of an experimental model of RMD during P10-20 for exploring anxiety and depression caused by child neglect and abuse.

## Supporting Information

**S1 Figure**

**No effect of stimulation on neuronal activity in the BLA.** (**A**) Zif268/Egr1 cells with no sRS are shown in the BLA. (**B**) Those with sRS are shown. (**C**) Their numbers are compared between the groups (Student’s t-test, *p* = 0.85). Error bars represent SEMs. sRS, single exposure to restraint stress. BLA, basolateral amygdala. Scale bar = 100 *µ*m.

**S2 Figure**

**GAD2-positive neurons contain GABA.** (**A**) GAD2 reporter-positive cells (magenta) are labeled with GABA immunostaining (green) in layer VI of the cerebral cortex. (**B**) A double-positive cell is shown in the LHb. Arrowheads indicate double-positive cells. GAD2, glutamic acid decarboxylase 2. Scale bar = 100 *µ*m.

**S3 Figure**

**No effects of RMDsp on perineuronal nets (PNNs) in the LHb.** (**A**) The PNNs are shown with WFA lectin-staining in the control (Aa) and RMDsp groups (Ab); the WFA-positive areas are compared between the groups (*p* = 0.58, Student’s t-test) (Ac). (**B**) The PNNs are shown with the aggrecan immunostaining for the control (Ba) and RMDsp (Bb) groups; their aggrecan-positive areas are compared between groups (*p* = 0.80, Student’s t-test) (Bc). Error bars represent SEMs. WFA, wisteria floribunda agglutinin; RMDsp, repeated maternal deprivation during the second part of early-life (P10-20). Scale bar = 100 *µ*m.

## Acknowledgments

We thank Drs. NISHIJO H, NISHIMARU H, and UEMATSU A for critical reading of the manuscript and Drs. TAKEUCHI Y and KAWAGUCHI M for discussions.

## References

1. T. Abe, H. Kiyonari, G. Shioi, K.-I. Inoue, K. Nakao, S. Aizawa, and T. Fujimori. Establishment of conditional reporter mouse lines at rosa26 locus for live cell imaging. genesis, 49(7):579–590, 2011.

2. B. Aisa, R. Tordera, B. Lasheras, J. Del Río, and M. J. Ramírez. Cognitive impairment associated to hpa axis hyperactivity after maternal separation in rats. Psychoneuroendocrinology, 32(3):256–266, 2007.

3. H. Aizawa, R. Amo, and H. Okamoto. Phylogeny and ontogeny of the habenular structure. Frontiers in neuroscience, 5:138, 2011.

4. P. L. Brown, H. Palacorolla, D. Brady, K. Riegger, G. I. Elmer, and P. D. Shepard. Habenula-induced inhibition of midbrain dopamine neurons is diminished by lesions of the rostromedial tegmental nucleus. Journal of Neuroscience, 37(1):217–225, 2017.

5. J. F. Enwright, S. Sanapala, A. Foglio, R. Berry, K. N. Fish, and D. A. Lewis. Reduced labeling of parvalbumin neurons and perineuronal nets in the dorsolateral prefrontal cortex of subjects with schizophrenia. Neuropsychopharmacology, 41(9):2206–2214, 2016.

6. S. Geisler, K. H. Andres, and R. W. Veh. Morphologic and cytochemical criteria for the identification and delineation of individual subnuclei within the lateral habenular complex of the rat. Journal of Comparative Neurology, 458(1):78–97, 2003.

7. J. A. Gordon and M. P. Stryker. Experience-dependent plasticity of binocular responses in the primary visual cortex of the mouse. Journal of Neuroscience, 16(10):3274–3286, 1996.

8. O. Hikosaka, S. R. Sesack, L. Lecourtier, and P. D. Shepard. Habenula: crossroad between the basal ganglia and the limbic system. Journal of Neuroscience, 28(46):11825–11829, 2008.

9. H. Ichijo, M. Hamada, S. Takahashi, M. Kobayashi, T. Nagai, T. Toyama, and M. Kawaguchi. Lateralization, maturation, and anteroposterior topography in the lateral habenula revealed by zif268/egr1 immunoreactivity and labeling history of neuronal activity. Neuroscience research, 95:27–37, 2015.

10. S. Kang, J. Li, W. Zuo, R. Fu, D. Gregor, K. Krnjevic, A. Bekker, and J.-H. Ye. Ethanol withdrawal drives anxiety-related behaviors by reducing m-type potassium channel activity in the lateral habenula. Neuropsychopharmacology, 42(9):1813–1824, 2017.

11. A. Kisner, J. E. Slocomb, S. Sarsfield, M. L. Zuccoli, J. Siemian, J. F. Gupta, A. Kumar, and Y. Aponte. Electrophysiological properties and projections of lateral hypothalamic parvalbumin positive neurons. PloS one, 13(6), 2018.

12. D. Knowland, V. Lilascharoen, C. P. Pacia, S. Shin, E. H.-J. Wang, and B. K. Lim. Distinct ventral pallidal neural populations mediate separate symptoms of depression. Cell, 170(2):284–297, 2017.

13. B. Li, J. Piriz, M. Mirrione, C. Chung, C. D. Proulx, D. Schulz, F. Henn, and R. Malinow. Synaptic potentiation onto habenula neurons in the learned helplessness model of depression. Nature, 470(7335):535–539, 2011.

14. K. Li, T. Zhou, L. Liao, Z. Yang, C. Wong, F. Henn, R. Malinow, J. R. Yates, and H. Hu. βcamkii in lateral habenula mediates core symptoms of depression. science, 341(6149):1016–1020, 2013.

15. M. Li, C. D’arcy, and X. Meng. Maltreatment in childhood substantially increases the risk of adult depression and anxiety in prospective cohort studies: systematic review, meta-analysis, and proportional attributable fractions. Psychological medicine, 46(4):717–730, 2016.

16. G. M. MacQueen, K. Ramakrishnan, R. Ratnasingan, B. Chen, and L. T. Young. Desipramine treatment reduces the long-term behavioural and neurochemical sequelae of early-life maternal separation. International Journal of Neuropsychopharmacology, 6(4):391–396, 2003.

17. R. A. Millstein and A. Holmes. Effects of repeated maternal separation on anxiety-and depression-related phenotypes in different mouse strains. Neuroscience & Biobehavioral Reviews, 31(1):3–17, 2007.

18. A. Minelli, N. Brecha, C. Karschin, S. DeBiasi, and F. Conti. Gat-1, a high-affinity gaba plasma membrane transporter, is localized to neurons and astroglia in the cerebral cortex. Journal of Neuroscience, 15(11):7734–7746, 1995.

19. S. Miyata, Y. Komatsu, Y. Yoshimura, C. Taya, and H. Kitagawa. Persistent cortical plasticity by upregulation of chondroitin 6-sulfation. Nature neuroscience, 15(3):414, 2012.

20. S. J. Mizumori and P. M. Baker. The lateral habenula and adaptive behaviors. Trends in neurosciences, 40(8):481–493, 2017.

21. J. Molet, P. M. Maras, S. Avishai-Eliner, and T. Z. Baram. Naturalistic rodent models of chronic early-life stress. Developmental psychobiology, 56(8):1675–1688, 2014.

22. J. Morris, K. Smith, P. Cowen, K. Friston, and R. J. Dolan. Covariation of activity in habenula and dorsal raphe nuclei following tryptophan depletion. Neuroimage, 10(2):163–172, 1999.

23. S. Ohayon, O. Avni, A. L. Taylor, P. Perona, and S. R. Egnor. Automated multi-day tracking of marked mice for the analysis of social behaviour. Journal of neuroscience methods, 219(1):10–19, 2013.

24. N. Omelchenko, R. Bell, and S. R. Sesack. Lateral habenula projections to the rat ventral tegmental area: Sparse synapses observed onto dopamine and gaba neurons. The European journal of neuroscience, 30(7):1239, 2009.

25. G. Paxinos and K. B. Franklin. Paxinos and Franklin’s the mouse brain in stereotaxic coordinates. Academic press, 2004.

26. G. Pietrini, Y. J. Suh, L. Edelmann, G. Rudnick, and M. J. Caplan. The axonal gamma-aminobutyric acid transporter gat-1 is sorted to the apical membranes of polarized epithelial cells. Journal of Biological Chemistry, 269(6):4668–4674, 1994.

27. C. D. Proulx, O. Hikosaka, and R. Malinow. Reward processing by the lateral habenula in normal and depressive behaviors. Nature neuroscience, 17(9):1146, 2014.

28. W. Rehan, J. Antfolk, A. Johansson, P. Jern, and P. Santtila. Experiences of severe childhood maltreatment, depression, anxiety and alcohol abuse among adults in finland. PLoS one, 12(5), 2017.

29. C. E. Ribak, W. M. Tong, and N. C. Brecha. Gaba plasma membrane transporters, gat-1 and gat-3, display different distributions in the rat hippocampus. Journal of Comparative Neurology, 367(4):595–606, 1996.

30. A. Ribic, M. C. Crair, and T. Biederer. Synapse-selective control of cortical maturation and plasticity by parvalbumin-autonomous action of syncam 1. Cell reports, 26(2):381–393, 2019.

31. J.-S. Seo, P. Zhong, A. Liu, Z. Yan, and P. Greengard. Elevation of p11 in lateral habenula mediates depression-like behavior. Molecular psychiatry, 23(5):1113–1119, 2018.

32. S. Sugiyama, A. A. Di Nardo, S. Aizawa, I. Matsuo, M. Volovitch, A. Prochiantz, and T. K. Hensch. Experience-dependent transfer of otx2 homeoprotein into the visual cortex activates postnatal plasticity. Cell, 134(3):508–520, 2008.

33. N. Tamamaki, Y. Yanagawa, R. Tomioka, J.-I. Miyazaki, K. Obata, and T. Kaneko. Green fluorescent protein expression and colocalization with calretinin, parvalbumin, and somatostatin in the gad67-gfp knock-in mouse. Journal of Comparative Neurology, 467(1):60–79, 2003.

34. H. Taniguchi, M. He, P. Wu, S. Kim, R. Paik, K. Sugino, D. Kvitsani, Y. Fu, J. Lu, Y. Lin, et al. A resource of cre driver lines for genetic targeting of gabaergic neurons in cerebral cortex. Neuron, 71(6):995–1013, 2011.

35. A. Tchenio, S. Lecca, K. Valentinova, and M. Mameli. Limiting habenular hyperactivity ameliorates maternal separation-driven depressive-like symptoms. Nature communications, 8(1):1–8, 2017.

36. C. Winter, B. Vollmayr, A. Djodari-Irani, J. Klein, and A. Sartorius. Pharmacological inhibition of the lateral habenula improves depressive-like behavior in an animal model of treatment resistant depression. Behavioural brain research, 216(1):463–465, 2011.

37. J. Yamada, T. Ohgomori, and S. Jinno. Perineuronal nets affect parvalbumin expression in gaba ergic neurons of the mouse hippocampus. European Journal of Neuroscience, 41(3):368–378, 2015.

38. L.-M. Yang, B. Hu, Y.-H. Xia, B.-L. Zhang, and H. Zhao. Lateral habenula lesions improve the behavioral response in depressed rats via increasing the serotonin level in dorsal raphe nucleus. Behavioural brain research, 188(1):84–90, 2008.

39. Y. Yang, Y. Cui, K. Sang, Y. Dong, Z. Ni, S. Ma, and H. Hu. Ketamine blocks bursting in the lateral habenula to rapidly relieve depression. Nature, 554(7692):317–322, 2018b.

40. Y. Yang, H. Wang, J. Hu, and H. Hu. Lateral habenula in the pathophysiology of depression. Current opinion in neurobiology, 48:90–96, 2018a.

41. L. Zhou, M.-Z. Liu, Q. Li, J. Deng, D. Mu, and Y.-G. Sun. Organization of functional long-range circuits controlling the activity of serotonergic neurons in the dorsal raphe nucleus. Cell reports, 18(12):3018–3032, 2017.

