## Supplementary figures and images for "Maturation of lateral habenula and early-life experience-dependent alteration with behavioral disorders in adulthood"

### S1 Figure

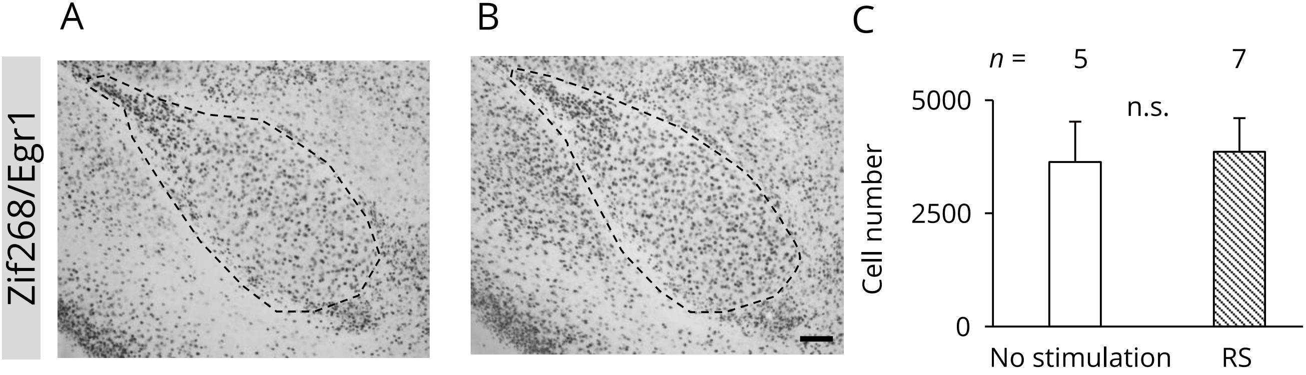

### S2 Figure

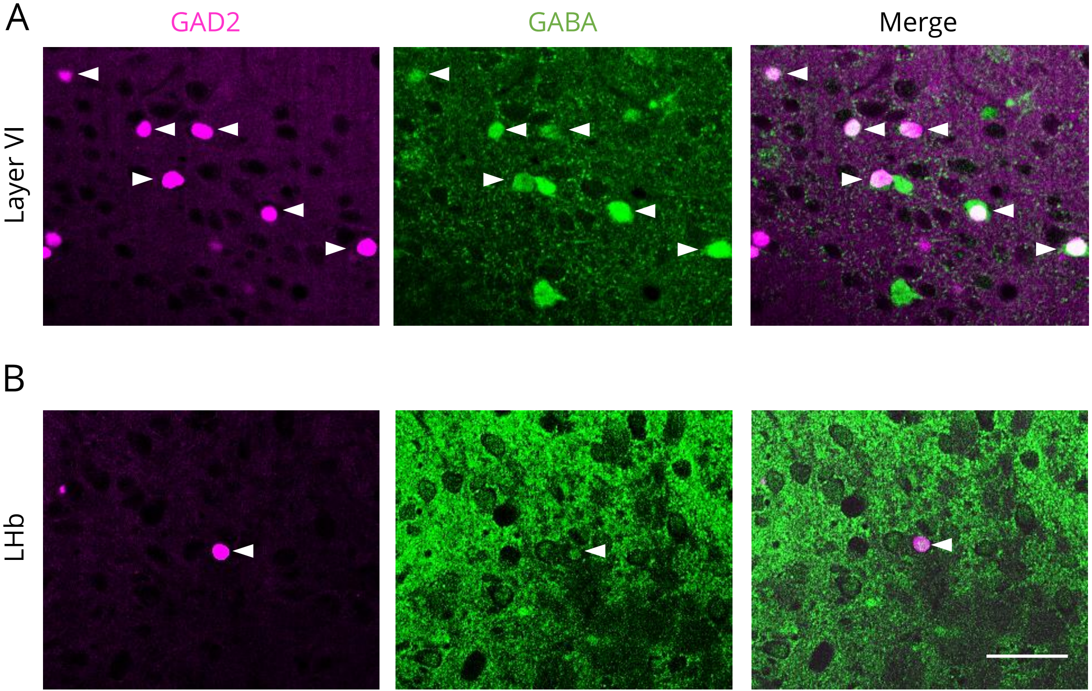


S2 Figure

### S3 Figure

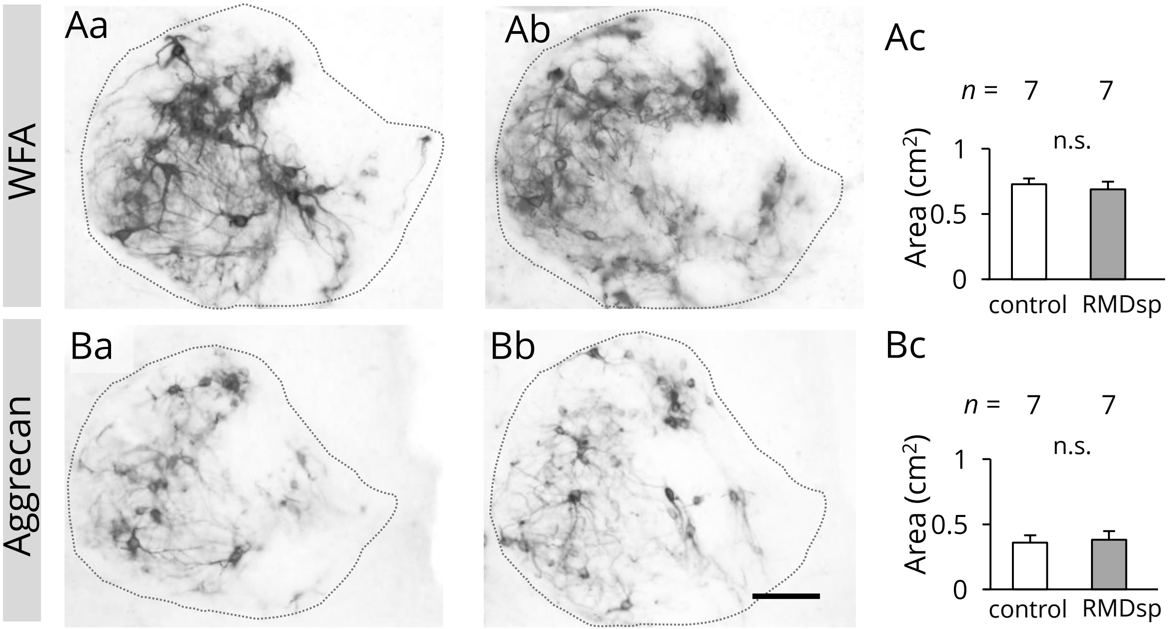
